# Analysis of sperm processing and cryopreservation procedures in the common marmoset (*Callithrix jacchus*) to reduce fragmentation and maximize total and progressive motility

**DOI:** 10.64898/2026.10.29.762870

**Authors:** Rebecca A. Holton, Joshua T. Brennan, Sophia I. Lipari, James Pickel, Nia Denney, Bhuchitra Singh, Jinwoo Jun, Yang Zhang, Pan P. Li, Colin D. Robertson, Ingie Hong, Rick Huganir, Reza Kalhor, Christopher A. Ross, James H. Segars, Xiaoqin Wang

## Abstract

The marmoset has emaserged as an ideal species for studying complex neurodevelopmental diseases, and IVF is required for gene-editing success. Reliable sperm processing is paramount to several IVF techniques, including traditional insemination. If not used immediately, thermal stress from cryopreservation diminishes sperm quality, requiring improved processing methods. Swim-up and gradient methods may limit adequate sperm quality, and the lack of a standardized freezing protocol often results in impaired post-thaw motility. Thus, reliable standards for sperm processing and cryopreservation in the marmoset are needed to support vital IVF processes. Collected semen underwent one of four pre-processing incubation steps before processing by swim-up, sucrose gradient, or a microfluidics assay. Sperm parameters were assessed using a Hamilton Thorne Computer Assisted Sperm Analysis (CASA) system. To evaluate cryopreservation, microfluidics-processed sperm were subjected to five protocols using different cryoprotectants and cooling methods. After thawing, sperm samples were analyzed with the CASA system and a DNA fragmentation assay. Incubation for 30 minutes at 38°C in TYH media decreased sperm aggregation, and subsequent microfluidics resulted in a higher percentage of motile and progressively motile sperm than swim-up or gradient methods. Our optimized cryopreservation protocol consistently achieved at least 20% progressive motility, a key benchmark for usable marmoset sperm, while previously published or commercial methods were less consistent. The cryopreservation protocol outperformed other published methods, having the least significant change in post-thaw motility without causing DNA damage. These findings establish standardized methods for reliably procuring and storing sperm samples for IVF or husbandry in the common marmoset.

## Introduction

Sperm processing and cryopreservation techniques are routinely used in mammalian reproductive research; however, there are no universally accepted protocols for the common marmoset (*Callithrix jacchus*). The common marmoset has become an important nonhuman primate model due to its small body size, early sexual maturity, short gestation period, and frequent twinning, which allow research colonies to be established and maintained more efficiently than those of larger primates [1]. Its physiological and neurological similarities to humans have also made it particularly useful in neuroscience, aging, and disease research [1]. More recently, the successful generation of transgenic marmosets has increased interest in assisted reproductive technologies (ART) to support the development and maintenance of genetically modified research models [2–3].

Reliable sperm processing methods are essential for ART, although the requirements for sperm quality and quantity differ depending on the procedure. Intracytoplasmic sperm injection (ICSI) requires only a small number of viable spermatozoa, whereas intrauterine insemination (IUI) requires sufficient numbers of progressively motile sperm. Conventional in vitro fertilization (IVF) additionally requires spermatozoa capable of capacitation and fertilization [4]. Sperm motility, DNA integrity, plasma membrane integrity, and acrosomal integrity are commonly used to assess sperm quality, as these characteristics can influence fertilization success and subsequent embryo development [5]. Processing methods must therefore recover enough functional spermatozoa while limiting damage associated with sample handling and preparation.

Sperm processing in the common marmoset is complicated by several characteristics of its ejaculate. Semen samples are typically small in volume and coagulate following collection, making it difficult to isolate motile spermatozoa [6–7]. Differences in sperm physiology and seminal plasma composition also limit the use of protocols developed for humans or other nonhuman primates, such as macaques [8]. Currently, in the marmoset field, there is no standard for sperm separation. Common sperm separation methods, including swim-up and density gradient centrifugation, can improve the proportion of motile sperm but often result in substantial losses in total sperm recovery [6–7]. These losses are especially problematic when working with small-volume or partially coagulated samples, where the number of sperm available for processing may already be limited. As a result, conventional methods may not consistently provide sufficient high-quality spermatozoa for ART, making it important to identify processing approaches better suited to marmoset semen.

Cryopreservation presents an additional challenge. The ability to freeze and store sperm would reduce the need for repeated semen collection and allow valuable genetic material to be preserved for future use. However, spermatozoa are susceptible to damage during freezing and thawing due to changes in temperature and osmotic conditions, ice crystal formation, membrane disruption, and oxidative stress [7]. These effects can reduce post-thaw motility and viability and compromise DNA and acrosomal integrity [9]. Previous studies in common marmosets have examined different cryoprotectants, including glycerol, as well as extender compositions, cooling rates, and freezing methods [5, 7, 10]. Despite these efforts, post-thaw sperm quality and recovery remain variable, and no single protocol has been consistently effective. This variability limits the practical use of cryopreserved sperm for ART, particularly in colonies where the number of available breeding males is limited.

Improving sperm processing and cryopreservation methods would have several practical benefits for marmoset reproductive research. More efficient recovery of motile sperm could increase the number of ART procedures performed from a single ejaculate, while reliable cryopreservation would allow sperm from valuable animals to be stored and used over time. These approaches could also help maintain genetic diversity within research colonies and support the preservation of genetically modified lines. Reducing the need for repeated semen collections may further minimize animal handling and improve the efficiency of colony management. In addition, methods developed for the common marmoset may help inform sperm preservation strategies in other nonhuman primates.

Given the limitations of existing methods, further optimization of sperm processing and cryopreservation is needed to improve the recovery of motile, functional spermatozoa in the common marmoset. In this study, we evaluated sperm processing and cryopreservation approaches with a focus on total and progressive motility before and after cryopreservation. Our goal was to improve the number and quality of spermatozoa available for ART and increase the number of males whose semen could be used successfully in reproductive procedures.

## Methods

All animal procedures were conducted in accordance with protocols approved by the Institutional Animal Care and Use Committee (IACUC) (Protocol No. PR24M269).

Common marmosets were maintained in single or paired housing under controlled environmental conditions, including a 12-hour light/dark cycle and regulated temperature and humidity. In paired housing, males were socially housed with females to maintain normal social interactions. Animals received a standardized Callitrichid Diet 5LK6 (LabDiet), consisting of Vitamin D3 and stabilized Vitamin C and enrichment 3 times per week, including 1 fresh produce, 1 dried food, and 1 protein-rich food with ad libitum access to water. Male ages ranged from 6 to 7 years.

### General Sperm Collection

With the aid of an assistant, male marmosets were hand-restrained in dorsal recumbency. After wiping clean the genital area with 70% alcohol, a Ferticare 2.0 massager was turned on, and the scrotum was stimulated continuously until ejaculation. Collection often occurred within the first 30 seconds. If no collection was obtained after 2 minutes of continued stimulation, the massager was turned off and the animal allowed to rest for at least 1 minute before a second or third 2-minute stimulation began.

### Aggregation

Approximately half of the semen sample from each monkey was appropriated to either an empty microfuge tube or one containing 400 µl warmed TYH medium. After collection, 4 µL from each sample was immediately added to a 2-chamber MicroCell slide (Vitrolife) with 20 µ chamber depth and imaged. The remaining sample was incubated at 38C for 30 minutes before another 4 µl aliquot was added to a separate slide and imaged.

### Sperm Processing Comparison

Each semen sample was pipetted into a microfuge tube containing 1 mL warmed modified Krebs-Ringer bicarbonate TYH medium. After liquifying in TYH for 30 minutes at 38C, sperm samples from each monkey were evenly split into three groups and assigned to a processing method: swim up, sucrose gradient, or the microfluidics assay (Figure 1). To compare freeze and thaw methods while limiting variables, sperm samples processed via the microfluidics method were subjected to one of five different protocols including Origio Sperm Freezing medium, the Morrell protocol using 5% glycerol [11], the O’Brien protocol using 6% glycerol [12], the Ogonuku (Sasaki) protocol with 7.5% glycerol [13], and a protocol adapted from the NIH using just 3% glycerol.

**Figure 1:**
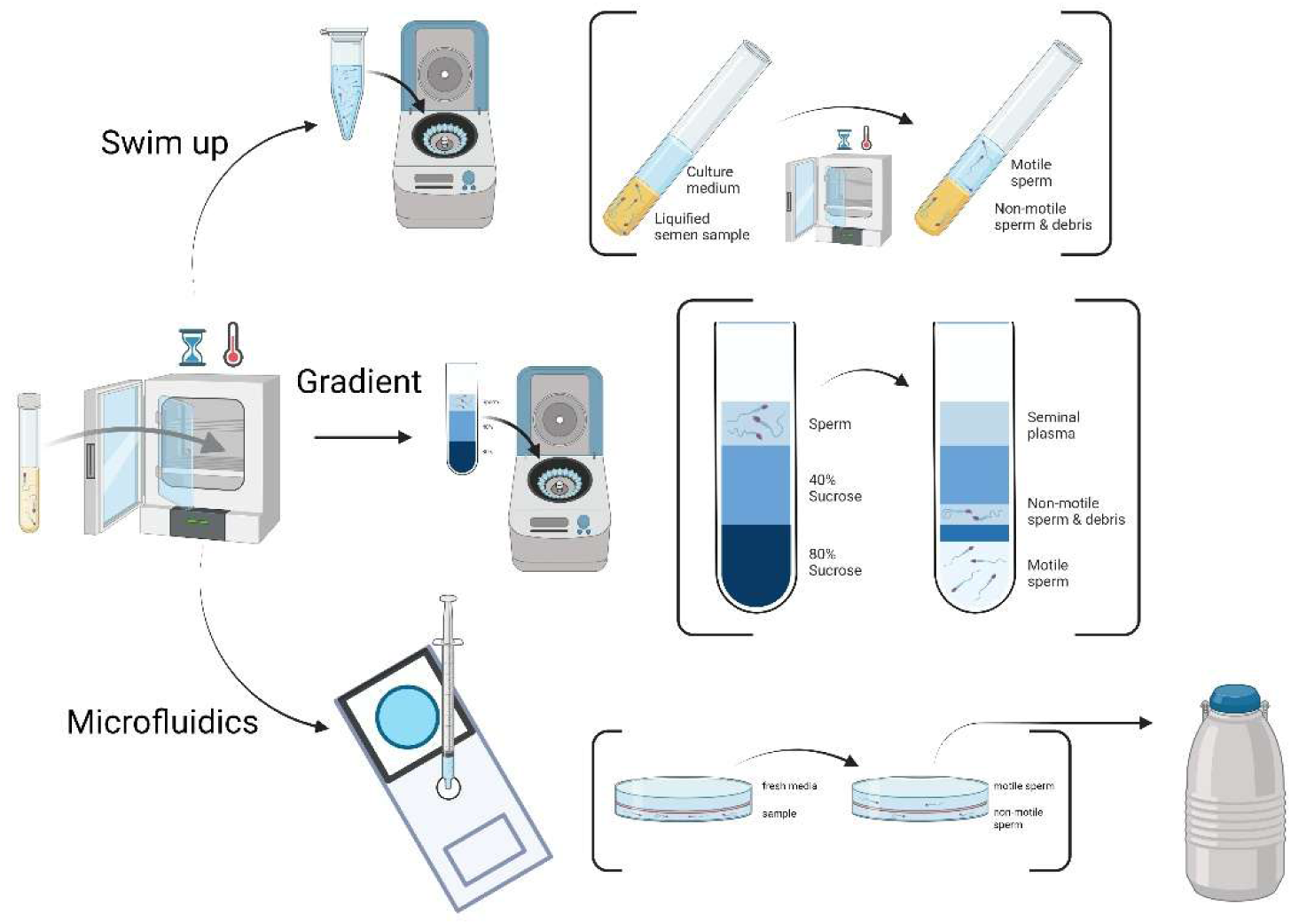
Comparison of Swim-up, Gradient, and microfluidics sperm processing methods. After incubation at 38C for 30 minutes, semen samples were processed using one of three methods to compare percentage of motile sperm and progressively motile sperm. Illustration created in BioRender. https://BioRender.com/8cnhgqd

A Hamilton Thorne CASA system was used to analyze sperm parameters and quantify sperm motility and progressive motility. Data collected were analyzed using an ordinary one-way ANOVA analysis followed by a Tukey’s multiple comparisons test, as well as the Student’s paired t-test with the p value corrected for multiple comparisons using the Holm-Šidák method.

### DNA Fragmentation

Microcentrifuge tubes containing agarose gel were warmed in a hot water bath (95-100C) for 90 seconds until the agarose melted. The microcentrifuge tubes were removed from heat and flicked to gather the droplets from along the tube wall and cap. After thawing, 30 µl of the semen sample was added to the tube and mixed with the agarose before immediately placing 30 µl of the mixture onto test wells 1 and 2. Glass coverslips were added and the slides were cooled at 2-8C for 5 minutes to solidify agarose. Slides were returned to room temperature, and the coverslip gently slid off to avoid damaging the gel.

Ten drops of DNA denaturant reagent were added dropwise to each test well, fully covering the slide. After leaving the slide at room temperature for 3 minutes, the slide was angled to allow excess liquid to drain off. For lysis, 10 drops of Lysis Solution were added to each test well and excess liquid similarly gently drained after 3 minutes of contact.

After adding 1 mL Distilled water and gently draining off excess liquid after 3 minutes, 50 µl of 75% Ethanol was added to each test well and the slides were immediately covered.

After the slides sat at room temperature for 1 minute, the covers were gently slid off and the slides gently rinsed with Distilled water to remove excess stain before air-drying at room temperature. Spermatozoa were observed under bright field microscopy (Figure 2) and the percentage of sperms with fragmented DNA was calculated with the below formulas for sperm DNA degradation and DNA Fragmentation Index (DFI):

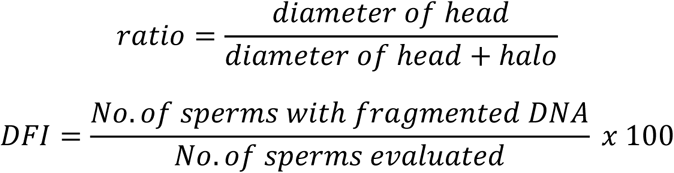

**Figure 2:**
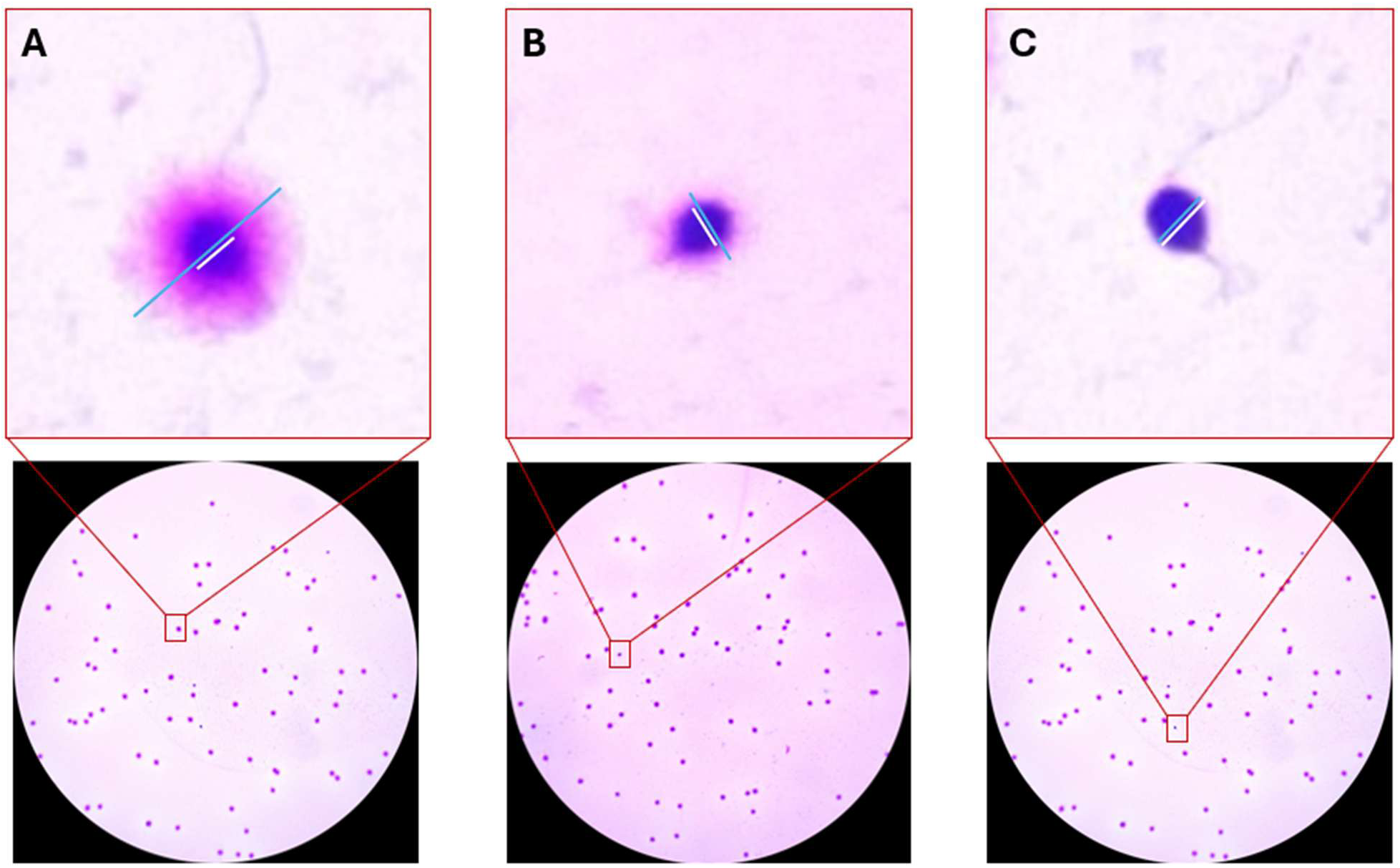
Measurement of sperm and halo. (A) Single spermatozoon with intact DNA. (B) Spermatozoon with partial DNA fragmentation, evidenced by the smaller pink halo. (C) Spermatozoon without a halo, indicating DNA fragmentation. (A-C) White lines indicate head diameter measurement. Blue lines indicate measurement of the combined head and halo. Lower images = 10x magnification; upper images enlarged to show detail.

After staining and completely drying, mounting medium was added to make permanent slides. Completed slides were stored at 2-8C.

Statistical analysis: All statistical tests were performed using GraphPad version 11. Sperm parameters between methods were analyzed using one-way ANOVA analysis followed by a Tukey’s multiple comparisons test. Changes pre- and post-thaw for individual samples were compared using a Student’s paired t-test with the p value corrected for multiple comparisons using the Holm-Šidák method. P-values <0.05 were considered statistically significant.

## Results

### Incubation and Media culture alleviates sperm aggregation

A leading challenge of marmoset sperm processing is the viscosity and commonality of aggregation of semen samples. Aggregation can greatly hinder sperm selection, regardless of the method. By comparing samples collected with and without media, and with and without 30 minutes of incubation at 38°C, we determined a method to liquify sperm samples prior to processing. Initial collection of a standard semen sample without the use of TYH media displayed large groups of sperm clumping together and forming raft-like forms with little movement (Figure 3A). In a particularly viscous sample, the aggregation was so severe that the sperm formed one large clump with varying density, rather than distinct clumps throughout the sample (Figure 3B). The addition of media immediately post collection allowed for the dilution and breakdown of these raft-like forms (Figure 3C). Even a more viscous sample showed improvement with the addition of TYH, displaying a more even density of sperm with less severe clumping (Figure 3D).

**Figure 3:**
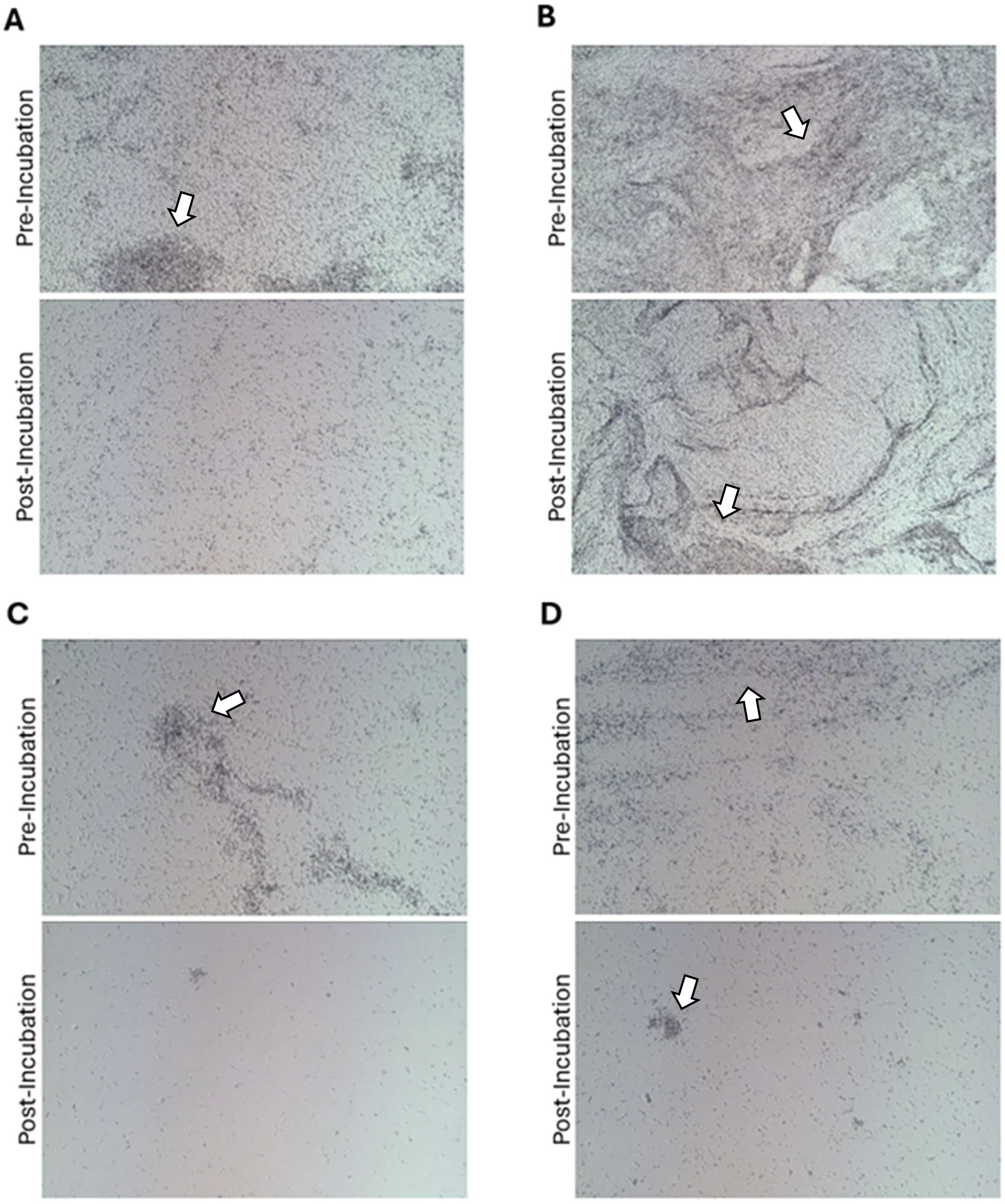
Incubation and the addition of media alleviated sperm aggregation in common marmoset semen samples. (A) An average semen sample from the common marmoset, without media, pre- and post-incubation. (B) A viscous semen sample from the common marmoset, without media, pre- and post-incubation. (C) An average semen sample with media pre- and post-incubation. (D) A viscous semen sample with media pre- and post-incubation. All images taken at 10x magnification. White arrow shows aggregation within specimens.

Samples, with and without media, were then incubated for 30 minutes, which further degraded the mucus and debris that commonly causes sperm aggregates to form. In the average sample, even without media, the clumps are less obvious (Figure 3A). The addition of media and incubation resulted in no aggregates at all, allowing for freely moving sperm (Figure 3C). A more viscous sample showed a similar pattern, with incubation loosening the sperm without media, with less density of aggregation, though clumping was still notable (Figure 3B, D). Incubation, combined with TYH resulted in a viscous sample having aggregates that were decreased in both size and number, increasing the amount of freely moving sperm in the sample. Overall, these results indicate that semen samples from marmosets may benefit from the addition of TYH media and incubation, allowing for easier sperm selection.

Microfluidic Sperm Separation Produced High-Quality Sperm Samples for IVF Though incubation and TYH media loosened aggregates, an appropriate IVF sperm sample will contain a high percentage of progressively motile sperm at high density, with few non motile sperm or other cellular debris. To achieve this, common forms of sperm preparation are: a swim-up assay, using the natural habit of sperm to swim up and towards the edges of a liquid to select for progressively motile individuals; a sucrose gradient separation, using centrifugation and varying solution density to separate out non-motile sperm and debris; and a microfluidics assay, which uses a thin, porous membrane separating the semen sample from fresh media, allowing motile sperm to swim up and into the fresh media.

With this experiment, we sought to identify the best means of sperm preparation to increase sperm sample quality. We began by comparing sample characteristics among each separation method by dividing each semen sample into three partitions and testing each partition with the three different methods of sperm preparation. The percentage of motile sperm in the sample before processing was 71.85%, with a concentration of 25.28 million motile sperm per milliliter (M/ml). Both the sucrose gradient and swim-up methods showed a statistically significant decrease in motile sperm, with average motility of 53.3% and 48.25%, respectively (p= 0.037, p=0.008, n=4 for each). Alternatively, the microfluidics assay showed the most significant change in motility; however, motility was significantly increased in post-processing samples, with 91.43% motility (p=0.027, n=4) (Figure 4A), and the concentration (6.15 M/mL) was comparable to that of the gradient and swim-up methods (6.625 M/mL and 4.36 M/mL, respectively). Microfluidics also had a significantly higher percentage of total motile sperm when compared to that of the gradient method (p=0.0002) and that of swim-up (p<0.0001).

**Figure 4.**
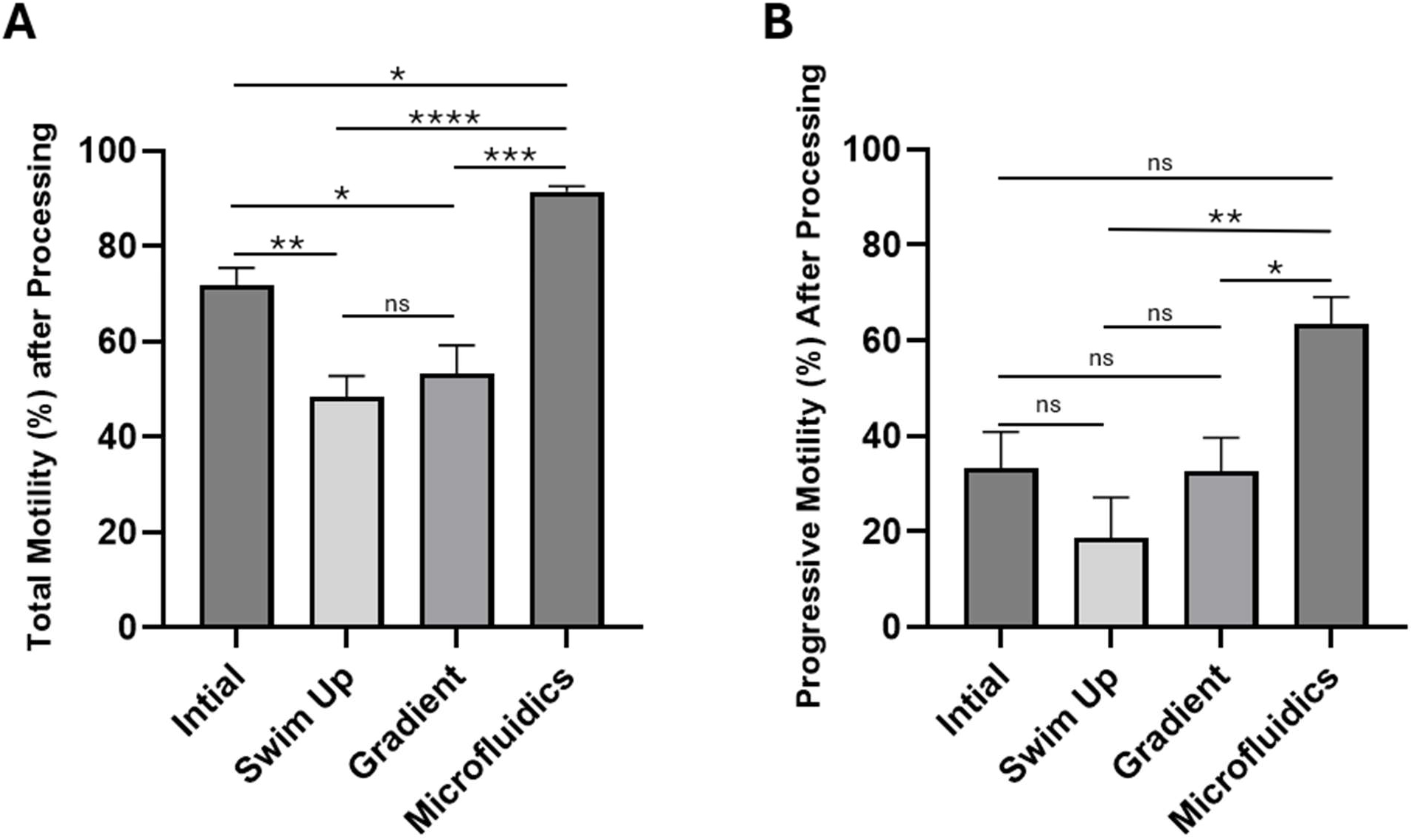
Microfluidic sperm processing improved the percentage of motile sperm and progressively motile sperm. (A) Microfluidics processing uniquely increased motile sperm percentage compared to initial mean (n=4, p=0.027). Swim-up and Gradient processing methods decreased motile sperm percentage, compared to initial mean (n=4, p=0.008 and n=4, p=0.037, respectively). (B) Microfluidics yielded the highest percentage of progressively motile sperm after processing (n=4, p=0.05, ns). Microfluidics showed a significant improvement in progressive motility when compared to swim-up (n=4, p=0.004) and gradient (n=4, p=0.045). A Hamilton Thorne CASA system was used to collect this data. All replicates were biologic replicates. Statistical significance determined by one-way ANOVA and Tukey’s multiple comparisons test. Error bars indicate standard error of the mean. Data represent a total of 4 biologic replicates, each with 3 technical replicates. NS=non-significant. * = p<0.05; ** = p<0.01.

We then compared the percent of sperm in each sample that was progressively motile, or with forward progression at a rapid pace. Our initial average was 33.33% progressive motility. The progressive motility of the sample processed with the sucrose gradient showed no change, with an average of 32.7% (p>0.99, n=4). The swim-up method also had no significant change in progressive motility, though the average was a lower 18.83% (p=0.54, n=4). The only method that showed a trend of improvement to progressive motility was the microfluidics assay, with a progressive motility of 63.53%, nearly double that of the initial sample (p=0.05, n=4) (Figure 4B). The microfluidics assay yielded more progressively motile sperm than either the gradient or swim-up methods (p=0.045 and p=0.0042, respectively). These data suggested that the microfluidics assay produced samples of sperm with the best parameters when compared with swim-up or gradient preparation methods.

### Evaluation of sperm cryopreservation methods

Transportation of marmoset sperm remains a challenge as there is not a universally accepted method of cryopreservation of marmoset sperm. For this reason, we tested several published methods of sperm cryopreservation to determine which protocol yielded the best results. We tested five methods referred to as follows: Sperm Freezing Medium (Origio), 5% glycerol [11], 6% glycerol [12], 7.5% glycerol [13], and the Hopkins protocol adapted from the NIH, referred to here as JHU. All samples were cryopreserved for at least 24 hours, after which they were thawed according to the JHU method as described elsewhere in this article. After thawing, we measured total motility and progressive motility. While all methods showed a significant decrease in both total and progressive motility post-thaw, the JHU method had the smallest reduction between total motility percentage before and after cryopreservation (53.37, Figure 5B, C). Notably, all methods still produced motile sperm post thaw, except the 6% glycerol method, in which all thawed sperm were motionless (Figure 5A). Next, we analyzed the percent of progressively motile sperm for each method. Once again, all methods showed a decrease in the percentage of progressive motility, but the JHU and Sasaki protocols were superior. The JHU method had the least significant change in progressive motility (31.37; p=0.06, Figure 5E, F). The mean of the NIH/Hopkins post-thaw samples (30.3%, Figure 5D) and was the only method that produced a sample with a value higher than the 30% progressive motility used as a benchmark for an adequate sperm sample.

**Figure 5:**
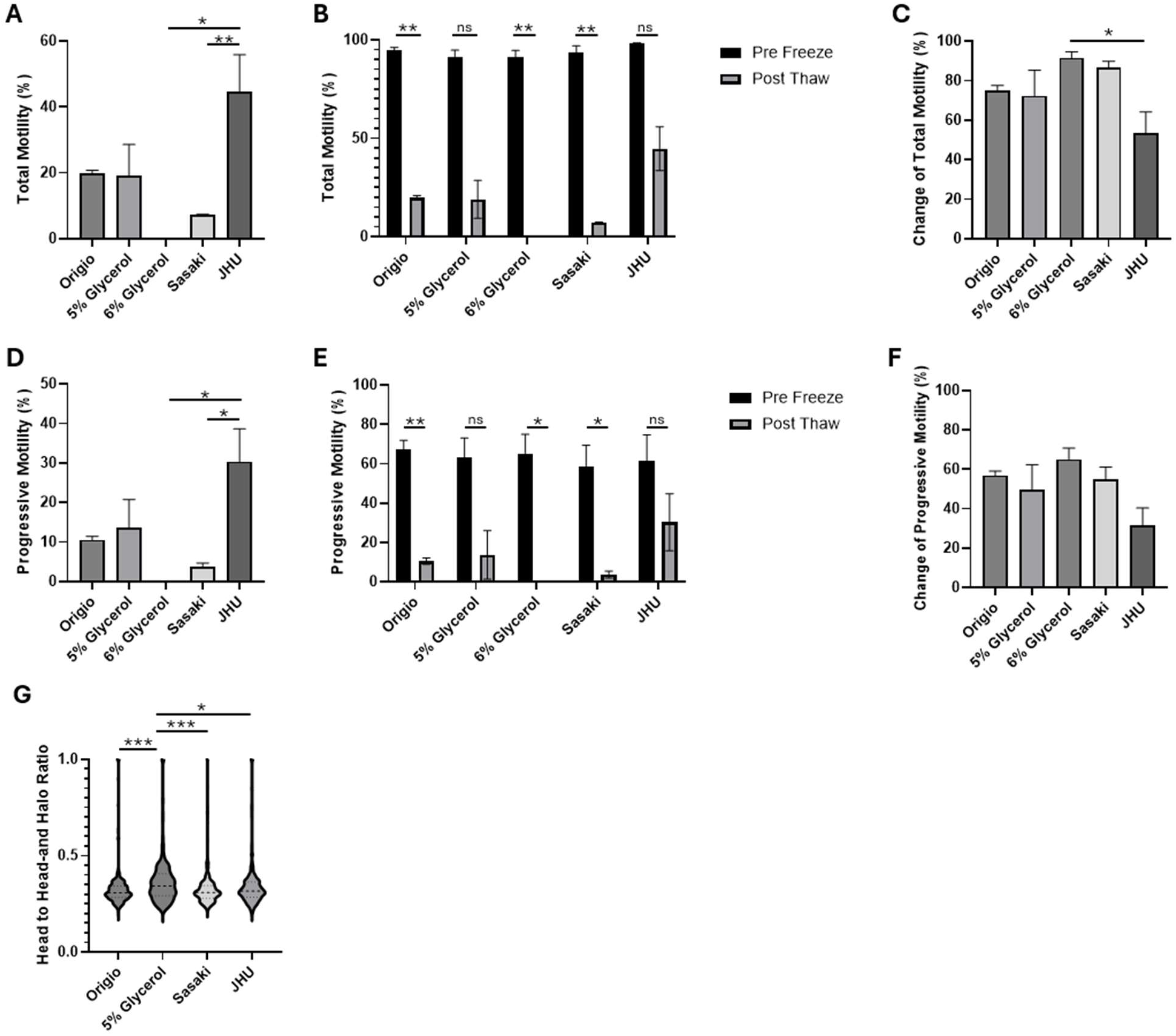
JHU method preserved motility and progressive motility after cryopreservation. (A) Percentage of total motility of sperm after cryopreservation and thawing. The JHU method showed a significant difference in comparison to the 6% glycerol method (p=0.005) and the Sasaki method (p=0.016). (B) The difference between percent total motility pre-cryopreservation and post-thaw. Origio (p=0.005), 6% glycerol (p=0.005), and Sasaki (p=0.005). (C) Post-thaw change in total motility. 6% glycerol showed a significantly larger change than JHU (p=0.04). (D) Percentage of progressive motility of sperm after cryopreservation and thawing. The JHU method showed more progressively motile sperm than the 6% glycerol (p=0.01) and Sasaki (p=0.02) methods. (E) The difference between percent progressive motility pre-cryopreservation and post-thaw: Origio (p=0.007), 6% glycerol (p=0.03), and Sasaki (p=0.03) all showed a significant decrease after thawing. (F) Post-thaw change in progressive motility. No significance was observed. (G) DNA fragmentation assay. The average head to head-and-halo ratio after cryopreservation. A smaller ratio indicated less DNA fragmentation. 5% glycerol had the highest ratio compared to the other methods. (A-F) Data represent a total of 4 biologic replicates, each with 3 technical replicates. Statistical significance determined by ordinary one-way ANOVA and Tukey’s multiple comparisons test (A, C, D, F, G). Student’s paired t-test with the p value corrected for multiple comparisons using the Holm-Šidák method (B, E). Error bars indicate standard error of the mean. (G) n=300 sperm per marmoset, 3 marmosets per group. * = p<0.05; ** = p<0.01.

Because cryopreservation induces cell stress, we next examined which of the cryopreservation methods produced samples motile sperm with the least DNA fragmentation. Using a DNA fragmentation assay, we measured the stained halos around each individual sperm (Figure 5G). We then calculated the ratio of the diameter of the sperm head to the diameter of the head and halo combined. A larger halo, and therefore a smaller head to head-and-halo ratio indicated less DNA fragmentation. Sperm with small or no halos were considered fragmented DNA positive, and therefore, of poor quality. None of the methods tested were outside the clinically excepted normal range (0-15%), with all methods having approximately 1% of all sperms testing positive for DNA fragmentation. The mean head to head-and-halo ratio was also somewhat even, as the JHU, Origio, and Sasaki methods all showed no statistical difference between one another (JHU vs Origio p=0.55, JHU vs Sasaki p=0.50, Origio vs Sasaki p=0.99). Only the 5% glycerol method had a slight increase in ratio value compared to the others (5% vs JHU p=0.046, 5% vs Origio p=0.0005, 5% vs Sasaki p=0.0004). Overall, none of the four tested methods damaged sperm DNA beyond the accepted normal range. In summary, the protocol described was satisfactory for processing and cryopreserving sperm using microfluidics assays and an accessible method for freezing.

## Discussion

The lack of a standard protocol for processing and cryopreservation of sperm in the marmoset field leads to vast differences in the effectiveness of methods to produce high quality and storable sperm samples. Here we aimed to compare the three common methods of sperm preparation, as well as compare four previously published methods used for sperm cryopreservation. The results of this study indicated that incubation and dilution in media greatly enhances semen processing of marmoset samples. Additionally, the best means of sperm preparation was a microfluidics separation device. We also determined a safe and accessible protocol for the cryopreservation of sperm that consistently yielded progressive sperm after thawing.

One of the hallmarks of a high-quality sperm sample is a high percentage of progressive motility. In humans, 30% progressive motility is considered the low threshold for a healthy and fertile sperm sample [14–15]. For a fresh sample, only the microfluidics assay yielded a progressive motility higher than 40%. In fact, this method showed an average progressive motility of 63.53%, higher than the average range of fertile humans as reported by the WHO (2021). The microfluidics method does not rely on correctly angling a tube, like swim-up, or the layering of sucrose solutions, like the gradient method, reducing the likelihood of human error. This method is consistent and beginner-friendly, making it ideal for academic laboratories.

Development of cryopreservation for marmoset sperm has been slow, with few published methods. One of the most effective methods recently presented [7] used a controlled rate freezer. This method, while effective, relies on equipment that is not readily available in many laboratories. Therefore, we sought to determine an alternative method that is more accessible and uses laboratory equipment already on site. By using only refrigeration and liquid nitrogen to cool the sperm, we propose a method that is less dependent on costly equipment that takes up valuable counter and floor space. When thawed, sperm cryopreserved with this method routinely yielded sperm with a higher progressive motility than 30%, indicating sperm that can successfully be used for IVF.

Optimizing these methods and increasing progressive motility in both fresh and cryopreserved samples can increase IVF success. Standardizing a protocol for marmoset sperm collection and storage will allow for more sharing of materials, shipment of sperm between colonies, and sperm banking of samples even after the male has died, as well as later use by labs internally. This will allow for more collaboration between laboratories globally, helping to increase the scope and breadth of NHP research.

## Authors’ Roles

XW, CAR, RLH, RK, and JHS acquired funding. RAH, JTB, JP, BS, JHS, and IH conceptualized and designed the studies. RAH, JTB, SIL, JP, ND, and JHS developed and performed experimental work and data analysis. All authors provided experimental and methodology feedback. RAH, JTB, SIL, JP, ND, BS, and JHS performed animal procedures. ND, YZ, and XW managed the marmoset population. XW, JHS, RLH, RK, and CAR supervised the study and design. RAH, JTB, SIL, and JHS wrote the manuscript. All authors contributed to the original manuscript or with editing and reviewing and approved of the final form.

## Acknowledgements

We would like to thank the veterinarian team, OR techs, and RAR staff at the Johns Hopkins University School of Medicine for their continued support with our marmoset population. We also thank the Johns Hopkins University School of Medicine Departments of Gynecology and Obstetrics, Biomedical Engineering, Neurobiology, and Neuroscience for their support and assistance with this work.

## Funding

This work was supported by National Institutes of Health U01DA056556 to XW, CAR, RLH, RK, and JHS and by the Brain Pool Plus (BP+) Program through the National Research Foundation of Korea (NRF) funded by the Ministry of Science and ICT (RS-2025-25403864, I.H.). JHS was supported, in part, by the Howard W. and Georgeanna Seegar Jones Endowment at the Johns Hopkins School of Medicine.

**Licensing:** CC-BY

## Competing Interests

The authors have no competing interests to disclose.

## Notes

### Competing Interest Statement

The authors have declared no competing interest.

